# Discovery of Fibrillar Adhesins across Bacterial Species

**DOI:** 10.1101/2020.12.07.414375

**Authors:** Vivian Monzon, Aleix Lafita, Alex Bateman

## Abstract

**Background:** Fibrillar adhesins are long multidomain proteins attached at the cell surface and composed of at least one adhesive domain and multiple tandemly repeated domains, which build an elongated stalk that projects the adhesive domain beyond the bacterial cell surface. They are an important yet understudied class of proteins that mediate interactions of bacteria with their environment. This study aims to characterize fibrillar adhesins in a wide range of bacterial phyla and to identify new fibrillar adhesin-like proteins to improve our understanding of host-bacteria interactions.

**Results:** By careful search for fibrillar adhesins in the literature and by computational analysis we identified 75 stalk domains and 24 adhesive domains. Based on the presence of these domains in the UniProt Reference Proteomes database, we identified and analysed 3,388 fibrillar adhesin-like proteins across species of the most common bacterial phyla. We found that the bacterial proteomes with the highest fraction of fibrillar adhesins include several known pathogens. We further enumerate the adhesive and stalk domain combinations found in nature and demonstrate that fibrillar adhesins have complex and variable domain architectures, which differ across species. By analysing the domain architecture of fibrillar adhesins we show that in Gram positive bacteria adhesive domains are mostly positioned at the N-terminus of the protein with the cell surface anchor at the C-terminus, while their positions are more variable in Gram negative bacteria. We provide an open repository of fibrillar adhesin-like proteins and domains to facilitate downstream studies of this class of bacterial surface proteins.

**Conclusion:** This study provides a domain-based characterization of fibrillar adhesins and demonstrates that they are widely found across the main bacterial phyla. We have discovered numerous novel fibrillar adhesins and improved the understanding of how pathogens might adhere to and subsequently invade into host cells.

## Background

Studying how bacteria interact with their host is essential for understanding bacterial infection processes and characterizing interactions with commensal bacteria. Adhesion to the host cell is the first step of a bacterial infection. Interfering in the adhesion process by Anti-adhesion therapies, mostly by stimulating a humoral immune response, can prevent infections and is therefore widely used for the development of vaccines, targeting host binding proteins [1]. Advanced vaccine candidates are for example targeting the binding domain of the M protein of Group A Streptococcus bacteria [2]. Fibrillar adhesins are a recently defined class of long and repetitive surface proteins, which are expressed by a variety of different bacteria [3]. Besides mediating the interaction between bacteria and host cells, fibrillar adhesins can be involved in biofilm formation and other cell-cell interactions [4, 5]. Their name is based on their filamentous structure, which arises from their characteristic protein architecture. This architecture is composed of tandemly repeated domains, that we name stalk domains for their function as elongated rodlike structures, and a domain with adhesive function (Figure 1). It is assumed that the stalk of tandem repeats projects the functional part of the protein, the adhesive domain, away from the bacterial cell surface and towards the attachment target, such as the host cell [6]. Few fibrillar adhesins have been studied so far and these come from a limited range of organisms. To our knowledge, no comprehensive review or study of fibrillar adhesins across bacterial species from a domain based perspective has been performed before. Fortunately, fibrillar adhesins appear to be composed of a relatively limited number of domain families, which allows their computational investigation. This study aims to create a comprehensive survey of fibrillar adhesins across bacterial phyla to gain understanding of their prevalence, distribution and composition. This work uses Pfam domain and motif composition to identify and characterize new potential adhesins. We hope that this study will guide researchers to investigate interesting new fibrillar adhesins, leading to an improvement in our understanding of microbial interactions in health and disease.

**Figure 1:**
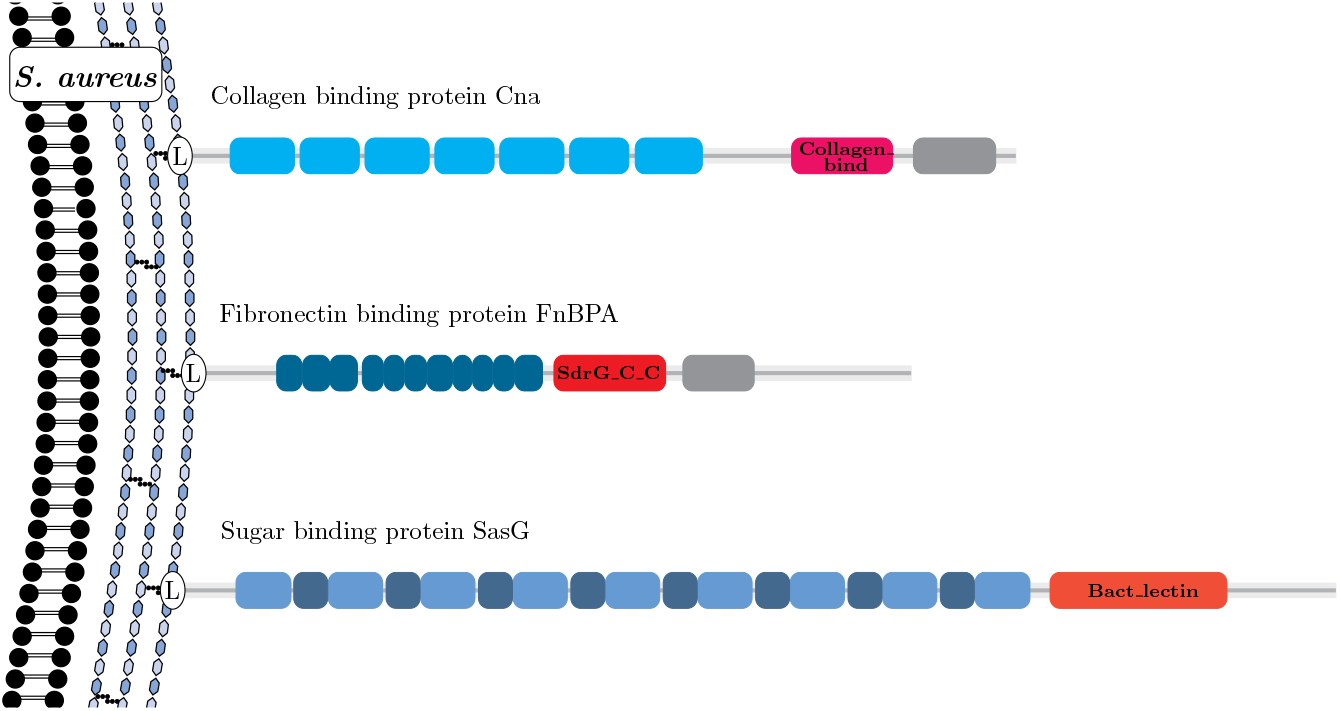
The domain architecture of fibrillar adhesins demonstrated by three well studied examples in *S. aureus*: The stalk domains (blue), adhesive domains (red) and host ligand targets (not shown) differ between the proteins. All three proteins are anchored to the peptidoglycan by the gram positive sortase anchor motif LPxTG, here indicated with the ‘L’.

## Results

### Detection of FA-like proteins

Fibrillar adhesins are composed of stalk domains, commonly repeated in tandem, and at least one adhesive domain. We gathered a set of 24 adhesive Pfam domain families from known fibrillar adhesins in the literature and 75 stalk domain families from proteins with tandem sequence repeats, as described in the Methods section. These 75 stalk domains include 6 new domain families, which were built in the course of this study (Pfam: PF19403 - PF19408). We used these domain families in our study to find further putative fibrillar adhesins, which we call fibrillar adhesin-like (FA-like) proteins.

The majority (15 out of 24) of the adhesive domains in our set bind to protein ligands, while 8 out of 24 bind to carbohydrates (supplementary table S2). The Ice_binding domain (Pfam: PF11999) differs from the other adhesive domains as it doesn’t bind to host cells or to other bacteria, but to ice crystals. It allows organisms to exist in extreme cold environments [7]. We identified in total 3,388 FA-like proteins composed of at least one adhesive and at least one stalk domain from our sets. These represent 0.012% of all proteins in the UniProt bacterial reference proteomes. Species-wise, FA-like proteins are found in 26% of the bacterial reference proteomes [8]. The large majority of these proteins are unstudied meaning we have identified numerous interesting targets for further investigation. Indeed, only 8.6% of the FA-like proteins are annotated with the GO term ‘cell adhesion’ in the UniProt database [8]. Some of these newly identified proteins are in well studied bacteria, for example we predicted three potential new internalin proteins (UniProt: Q8Y7I8, Q8Y8U2, Q8Y9W0) in *Listeria monocytogenes*. Different combinations of adhesive and stalk domains are found across FA-like proteins. The total number of identified stalk domains per protein varies from 1 to 73 in the FA-like proteins. Figure 2a shows the number of proteins identified based on the minimum number of stalk domains annotated in the FA-like protein. The largest number of FA-like proteins are found when only one known stalk domain is required in addition to the adhesive domain, and our study is based upon that set. Nevertheless, in 1,721 FA-like proteins three or more stalk domains were found in combination with an adhesive domain.

**Figure 2:**
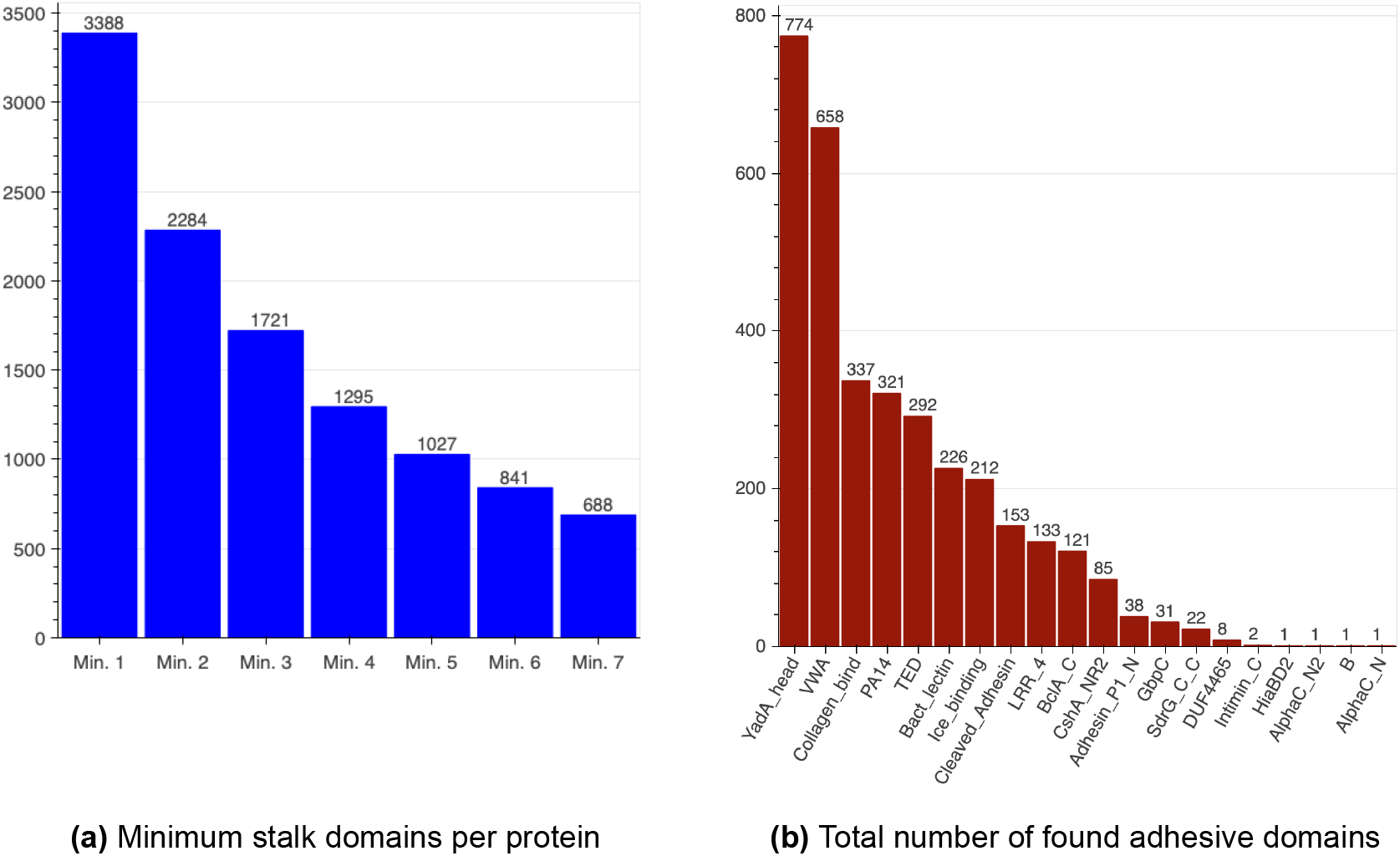
Minimum number of stalk domains per protein and the proportion of identified adhesive domains in FA-like proteins: a) Graph showing the number of FA-like proteins identified in our screen depending on the number of stalk domains required. Stalk domains can come from different stalk domain families. b) Graph showing the frequency of detected adhesive domains in combination with at least one known stalk domain. For proteins with multiple adhesive domains, each domain family is counted once per protein. Throughout this study we will use blue colouring for stalk domains and red colouring for adhesive domains.

### Prevalence of adhesive domains

Bacteria use a wide range of different protein domains to mediate adherence to their host or other organisms. To date it is not known which of these domains are the most common within fibrillar adhesins. Our domain based screen for FA-like proteins (Figure 2b) shows that the most commonly found adhesive domain in combination with at least one known stalk domain is YadA_head (Pfam: PF05658), found in 774 FA-like proteins (22.85 %), followed by VWA (Pfam: PF00092) with 658 hits (19.42 %) and Collagen_bind (Pfam: PF05737) found in 337 FA-like proteins (9.95 %).

### Domain combinations of FA-like proteins

We were further interested to learn if there is a common domain grammar in FA-like proteins. In the detected FA-like proteins combinations of 73 different stalk domains with 20 different adhesive domains were found. SlpA (Pfam: PF03217) and I-set (Pfam: PF07679) were the only stalk domains not found in combination with a known adhesive domain and the adhesive domains SabA_adhesion (Pfam: PF18304), SSURE (Pfam: PF11966), FadA (Pfam: PF09403) and FimH_man-bind (Pfam: PF09160) were not found with any known stalk domains. We plotted the combinations of adhesive and stalk domains as a heatmap, shown in figure 3. Some adhesive domains seem to be promiscuous and are found in combination with a wide range of different stalk domains, like the PA14 domain (Pfam: PF07691), whereas others are so far only found in combination with a specific stalk, like the AlphaC_N (Pfam: PF08829) which only appears in the combination with the Rib domain. Certain domains are mostly found together in a specific protein type, for example Adhesin_P1_N (Pfam: PF18652) with GbpC (Pfam: PF08363) in combination with the stalk domains Antigen_C (Pfam: PF16364), AgI_II_C2 (Pfam: PF17998) and Strep_SA_rep (Pfam: PF06696) as found in the *Streptococcus mutans* adhesin P1 protein [9].

**Figure 3:**
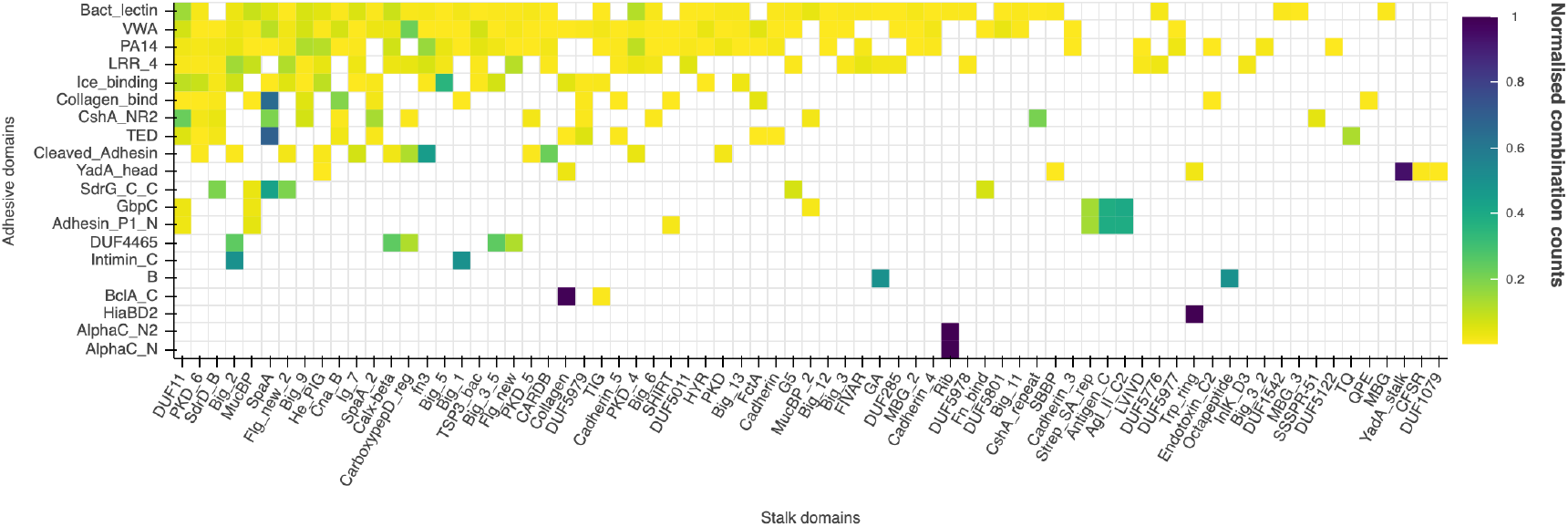
Stalk and adhesive domain combinations in FA-like proteins from the UniProt reference proteomes: The x-axis of this heatmap represents the known stalk domains and the y-axis the known adhesive domains. If the same domain family is found multiple times on the same protein, it was only counted once.

YadA_head with YadA_stalk (Pfam: PF05662) was the most prevalent domain combination found in 768 FA-like proteins. YadA_stalk was by far the most commonly occurring stalk do-main with annotations in 768 of the 3,388 FA-like proteins, followed by SpaA (Pfam: PF17802) with 668 FA-like protein hits (supplementary figure S2). Collagen_bind with SpaA was the second most prevalent domain combination found in 303 FA-like proteins.

It is important to keep in mind that there are probably more adhesive and stalk domains, which are not known yet and therefore not included in this study. Furthermore, only the UniProt reference proteomes were searched so that FA-like proteins and novel domain combinations can be missed, which are only found in the whole UniProtKB database. Moreover, imperfect annotation of stalk or adhesive domains by the Pfam HMMs may lead to missing annotations. One example is the SasY fibrillar adhesion of *S. aureus* with AlphaC_N and AlphaC_N2 adhesive domains in combination with Big_6 stalk domains. SasY is not found in the UniProt reference proteomes, but furthermore its adhesive domains are not annotated in Pfam.

Besides combinations of adhesive and stalk domains, which occur together in the same protein, there are also stalk domains that are commonly found together (supplementary figure S3). SpaA and Cna_B (Pfam: PF05738) are the stalk domains most often found together, in a total of 99 FA-like proteins. Both domain families belong to the Transthyretin clan (Pfam: CL0287), indicating that they are homologous and possess the same fold. SpaA is indeed found with a range of stalk domains, of which several belong to the Transthyretin clan, but others also belong to other clans, like Big_9 of the E-set clan (Pfam: CL0159). Of the 73 stalk domains which are found in combination with an adhesive domain, there are five domains, which are not found (in the Reference Proteomes) with another known stalk domain on the same protein. These five domains are Rib (Pfam: PF08428), QPE (Pfam: PF18874), Big_3 (Pfam: PF07523), Big_3_2 (Pfam: PF12245) and DUF5977 (Pfam: PF19404).

Domain pairs, which are not only annotated together in the same protein, but also function together, are called supra-domains [10]. In 224 out of the 337 FA-like proteins with Collagen_bind and in 20 of 22 FA-like proteins with SdrG_C_C (Pfam: PF10425), a Big_8 (Pfam: PF17961) domain was found N-terminal within 50 amino acid distance. The principles of the binding mechanisms of SdrG_C_C, called Dock lock latch (DLL) mechanism and of Collagen_bind, called Collagen-hug, are comparable. Both, in the DLL as well as in the Collagen-hug binding mechanism, two domains build a trench in which the ligand docks and is finally locked. By comparing the sequence annotation with the structure of the respective binding mechanism, it is clear that Big_8 is one of the two domains and so essential for the binding. Thus we define two supradomains as Big_8:SdrG_C_C and Big_8:Collagen_bind. Aligning the structures of both adhesive domains showed that they are structurally homologous with a RMSD < 5Å [11].

Another example of a supra-domain represented on the heatmap in figure 3 is the AlphaC_N and AlphaC_N2 domain, which form the N-terminal domain in combination with the Rib stalk domain repeats in the Alpha C protein in Group B Streptococcus [12].

### FA-like protein cell anchoring

As surface proteins, fibrillar adhesins have some sort of attachment to the bacterial cell surface. This can take many forms, such as being non-covalently attached by S-layer homology domains [13], or covalently anchored by a sortase anchoring motif. To better characterise FA-like proteins we also created a list of known anchoring motifs, signals and domains (supplementary table S3). The frequency of each of these different types of anchoring methods is shown in figure 4 below. For most of the detected FA-like proteins (2,038 in all phyla), none of the known anchor domains or motifs listed in supplementary Table S3 were found (labelled as ‘No match’). For those proteins lacking a known anchor we supposed they may be anchored with a transmembrane spanning helix. When searching with TMHMM we found transmembrane regions for only 10.6% of the FA-like proteins without known anchor domains. The Gram positive anchor motif is the most prevalent one in the detected FA-like proteins with 662 hits in total, of which 482 FA-like proteins belong to Firmicutes and 165 belong to the Actinobacteria phylum. In addition to the LPxTG motif the alternative Gram positive anchor motifs ‘LPxTA’, ‘NPxTG’, ‘LPxGA’, ‘LAxTG’ and ‘NPQTN’ were included [14]. The S-layer homology domain was found in 73 detected FA-like proteins and again mostly in the generally Gram positive Firmicutes. The anchor domain CHU_C is found in 134 FA-like proteins, all of which belong to the Bacteroidetes phylum. Por_Secre_tail is found in 118 FA-like proteins and mostly in Bacteroidetes. Four proteins with LysM domains (Pfam: PF01476) were detected, three proteins with Choline_bind_1 (Pfam: PF01473), two proteins with Choline_bind_3 (Pfam: PF19127), two with IAT_beta and one with GW domain. In Proteobacteria the YadA anchor (Pfam: PF03895) domain and the ‘LapG + HemolysinCabind’ anchor were the most prevalent. The LapG motif is the protease cleavage site at the N-terminus of RTX adhesins, where the proteins are cleaved by the LapG protease and subsequently inserted into a beta barrel of an outer membrane pore [15]. To be certain to find the LapG cleavage site in RTX adhesins, we additionally searched for the characteristic RTX repeats, HemolysinCabind domain (Pfam: PF00353), at the C-terminus, which is a T1SS signal [16, 17].

**Figure 4:**
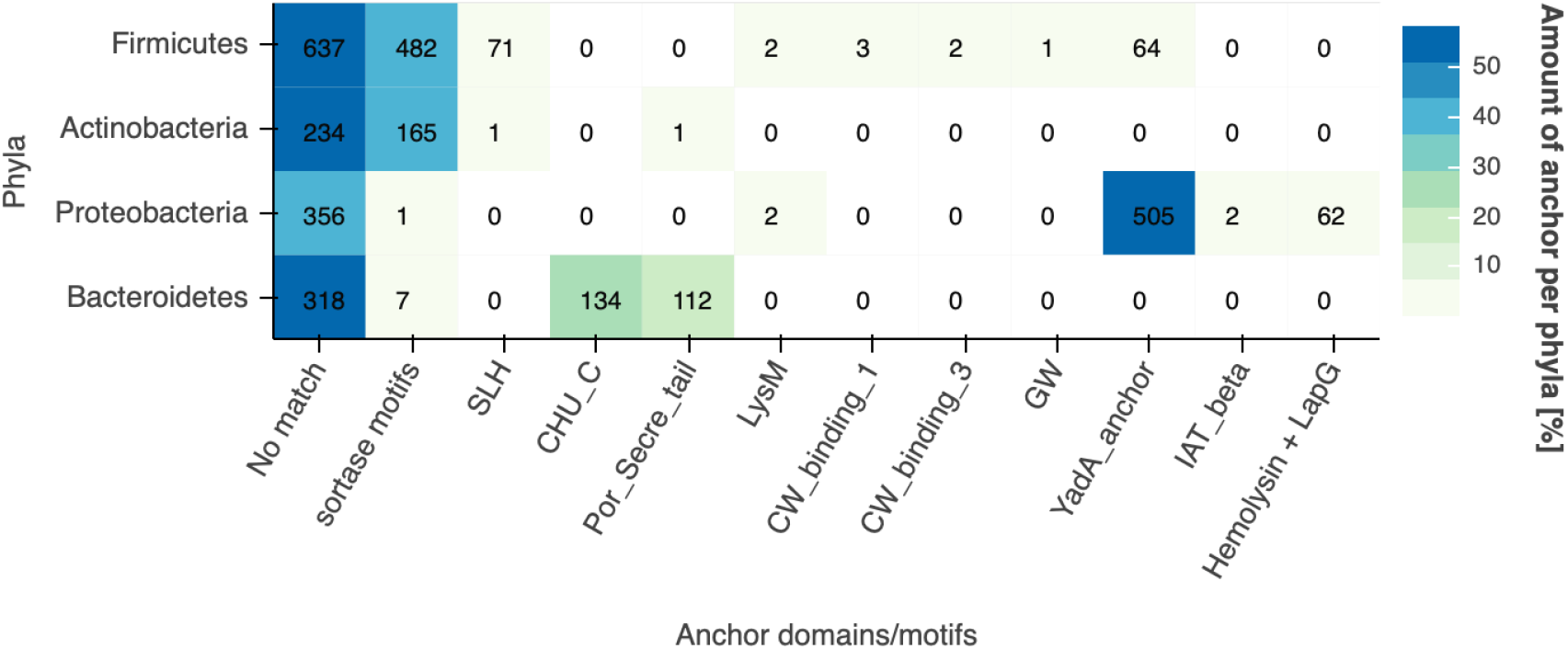
Prevalence of anchoring motifs and domains in FA-like proteins: The heatmap shows the anchor domains and motifs found in the FA-like proteins of the four main represented phyla. The colors represent the percentage abundance per phyla. The total number of anchors identified is found in each cell.

We ran PSORTb to identify that 860 of the 3,388 FA-like proteins were predicted to be localised at the cell wall, 529 to be at the outer membrane, 424 to be extracellular and 1348 of these proteins had no localisation predicted [18].

### Domain architecture of FA-like proteins

It is assumed that the adhesive domain is projected away from the bacterial cell surface by the elongated stalk. In this way its interaction with the host cell is promoted. This architecture is similar to multiprotein pili adhesins, where the protein with the binding function is the membrane distal one of all pili subunits [19]. In fibrillar adhesins the adhesive domain should be at the distal end of all stalk domains compared to the anchor. To investigate this assumption, the domain architectures of the identified FA-like proteins were investigated. In known fibrillar adhesins the adhesive domain is indeed often found at the N-terminus, far away from the cell wall anchoring motif at the C-terminus, e.g. in Cna of *S. aureus* [20, 21]. In order to see where in the protein the adhesive and stalk domains are annotated, their positions were plotted in figure 5. The plots are split between the most prevalent phyla of the detected FA-like proteins, which are Firmicutes (36.63%), Proteobacteria (27.33%), Bacteroidetes (16.74%) and Actinobacteria (11.81%). For Actinobacteria and Firmicutes we see a strong signal with the adhesive domain found at the N-terminus and the stalk domains found towards the C-terminus of the proteins (Figure 5a,b). This can be explained by the presence of the sortase anchor motif. The anchor motif is mostly found in Gram positive bacteria, where it gets anchored to the peptidoglycan cell wall. Interestingly we also see the same pattern in proteins lacking the sortase motif. This is most likely due to some sortase motifs being missed by current search methods. Interestingly, the FA-like proteins without a sortase motif in Firmicutes do show a small peak at the C-terminus, which might point out a different anchor method being used.

**Figure 5:**
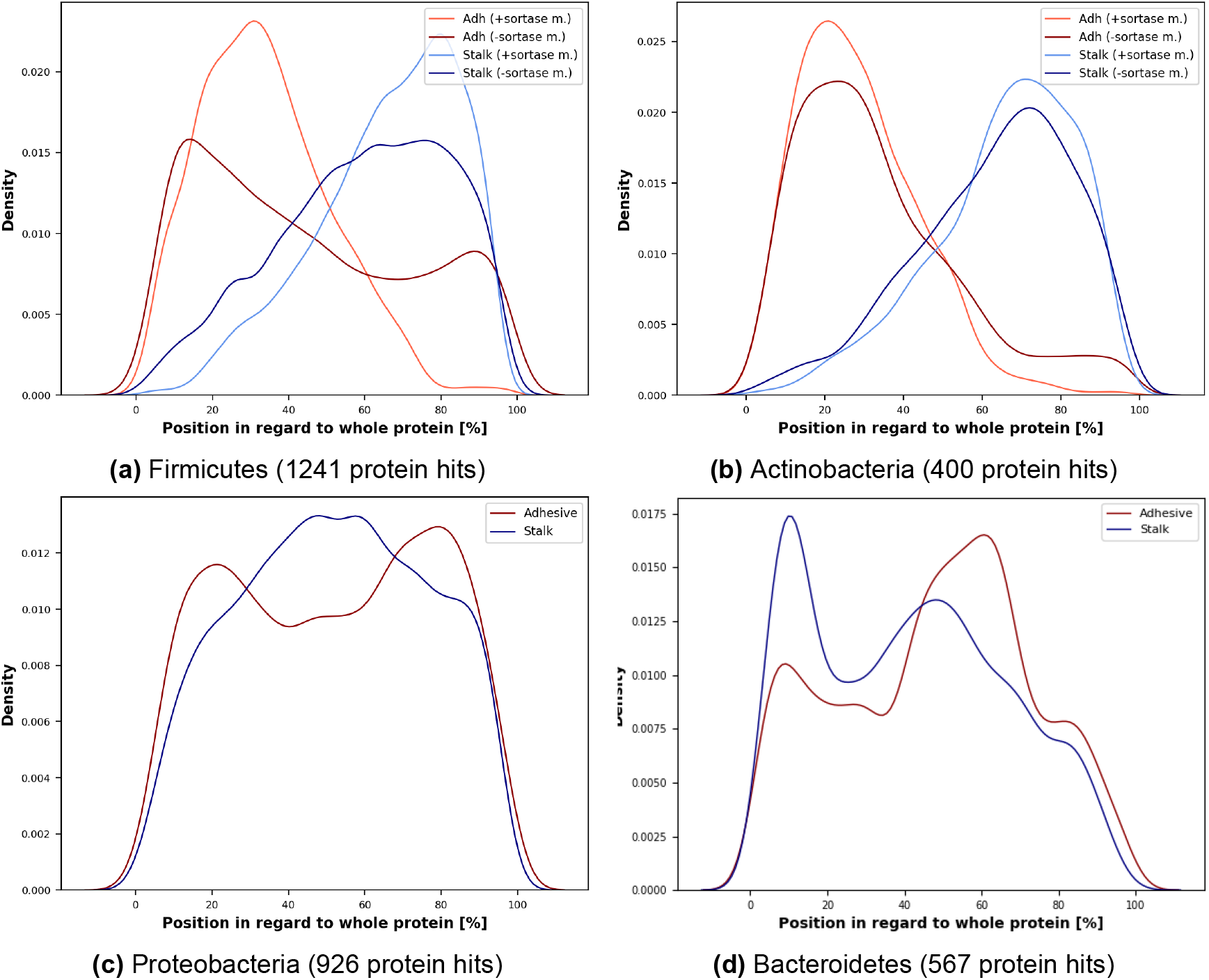
Domain architecture of FA-like proteins split by bacterial phyla: In these density plots the position of adhesive and stalk domains in FA-like proteins of the four main represented taxonomic phyla are plotted. 0% on the x-axes hereby represent the N-termini and 100% the C-termini of the proteins. The data are split between proteins with sortase anchor motif or without for the Gram positive Firmicutes and Actinobacteria phyla.

In the Gram negative phyla Bacteroidetes and Proteobacteria, the distribution of domains is more variable. For Proteobacteria (Figure 5c), the adhesive as well as the stalk domains show a relatively even distribution along the length of the FA-like proteins with a weak preference for adhesive domains at the termini. In Bacteroidetes, the stalk and adhesive domain distributions are similar, (Figure 5d) but surprisingly many adhesive domains are to be found centrally. There are known examples of centrally located adhesive domains. One example is adhesion P1 pro-tein in *S. mutans*, where the adhesive domain Adhesin_P1_N is annotated at the N-terminus and a second adhesive domain GbpC is annotated in the middle of the protein sequence. Ad-hesin_P1_N folds then to interact with the stalk domain at the C-terminus, which results in having GbpC at the top of the protein [9].

### Taxonomic overview

To get a deeper knowledge about the origins and evolution of FA-like proteins we looked at the distribution of adhesive and stalk domains across the bacterial phyla (see figure 6). The FA-like proteins we detected are widespread across the bacterial tree of life. FA-like proteins were found in 29 of 56 phyla represented in the UniProt reference proteomes. 14 of the phyla without FA-like proteins have only a single reference proteome, suggesting that the apparent lack of FA-like proteins may be due to lack of sampling of proteomes currently.

**Figure 6:**
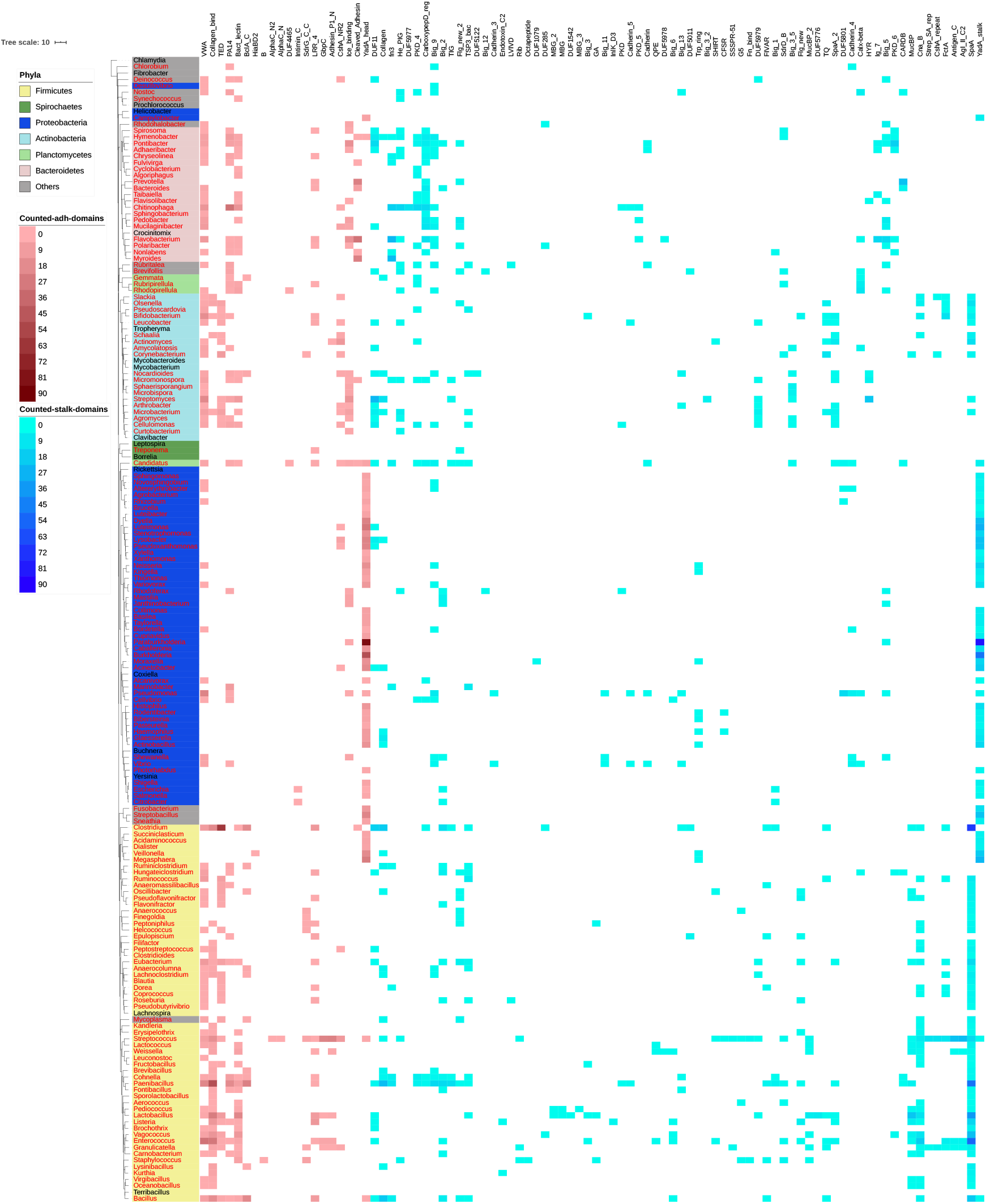
Taxonomic overview of adhesive and stalk domains in FA-like proteins: This taxonomic tree is built from representative bacterial genera of Firmicutes (yellow), Spirochaetes (dark green), Proteobacteria (dark blue), Actinobacteria (light blue), Planctomycetes (light green), Bacteroidetes (light red) and other phyla (grey). For the genera marked in red text FA-like proteins could be detected. To visualise which adhesive and stalk domains were mainly found in combination in the genera, they are mapped in blue and red respectively.

The adhesive and stalk domains in FA-like proteins mapped to the genera in figure 6 shows that overall the domain composition of Gram positive Firmicutes and Actinobacteria are similar to each other, but different from the Gram negative Proteobacteria. There are also Gram negative genera in Firmicutes, *Succiniclasticum*, *Acidaminococcus*, *Dialister*, *Veillonella* and *Megasphaera*, whose domain composition are more similar to the Proteobacteria. This underlines the importance of the cell surface composition for the architecture of FA-like proteins. In Proteobacteria the FA-like proteins are mainly built out of YadA_head and YadA_stalk domains. For nearly all of the Firmicutes genera and many Actinobacteria FA-like proteins composed of Collagen_bind, VWA or TED in combination with the SpaA stalk domain could be detected. The Gram negative Bacteroidetes use a wider range of domains compared to the Proteobacteria, of which many adhesive domains are the same as in Gram positive bacteria.

There are adhesive and stalk domains found in a wide range of genera, as well as adhesive and stalk domains limited to a few specific genera. VWA for example is an adhesive domain, which is widely distributed and which was found in combination with different stalk domains in Firmicutes, Proteobacteria, Bacteroidetes, Planctomycetes and Actinobacteria. On the other hand Intimin_C (Pfam: PF07979), seems to be adapted to Proteobacteria as it was only detected in this phylum in combination with the Big_1 (Pfam: PF02369) and Big_2 (Pfam: PF02368) stalk domains.

### Boundary between stalk and adhesive domain definitions

We observed that some of the domains regarded as adhesive in our list can also be found in repeats, leading us to wonder whether adhesive and stalk domain functions might be interchangeable. Multiple adhesive domains repeated in tandem might form stalks, and stalk domains might evolve adhesive functions over time to increase the binding avidity to host cells. In figure 7 we can see that most adhesive domains are only found once per protein. Exceptions are SSURE, B, YadA_head, Collagen_bind and LRR_4. Contrary to Collagen_bind and B domains, the SSURE domain is not found in combination with known stalk domains and we hypothesise that it may function as both adhesive and stalk domain. The Fn_bind (Pfam: PF02986) domain was defined as a stalk domain because it is found in combination with the adhesive domains SdrG_C_C and VWA. However, it is known that Fn_bind also has a binding function and can bind to Fibronectin, e.g. in the Fibronectin-binding proteins of *S. aureus* [22]. Suggesting that Fn_bind is another example of a dual adhesive and stalk domain.

**Figure 7:**
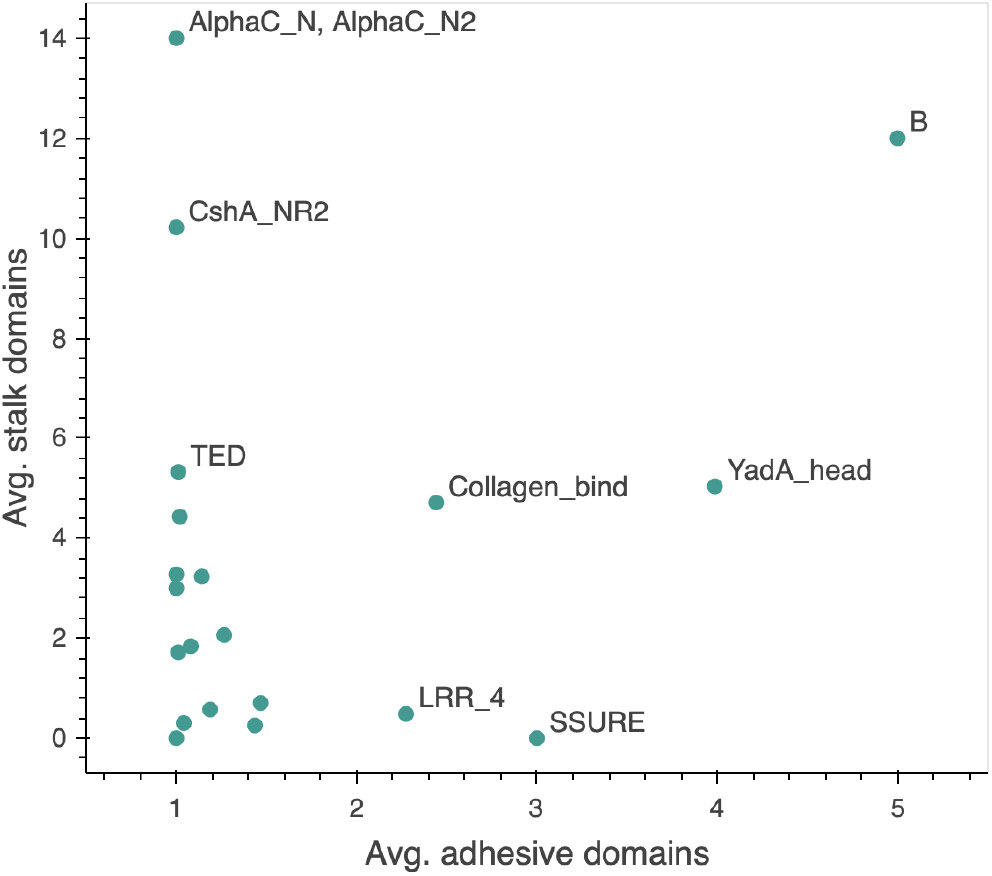
Average of domains found per adhesive domain family: For each adhesive domain family the average number of adhesive domains is plotted against the average number of stalk domains.

### Are FA-like proteins indicators of bacterial pathogenicity?

FA-like proteins can mediate pathogen-host interactions, but also interactions between bacteria, for example in biofilm formation. In the course of our study, we detected a correlation between pathogenicity and the presence of FA-like proteins for the well characterized organisms *Bacillus subtilis* and *Bacillus cereus*. For the non-pathogenic *Bacillus subtilis* no FA-like proteins were detected, compared to the pathogenic *Bacillus cereus* species, for which four different FA-like proteins were detected. Due to a lack of detailed annotation of pathogenic status of UniProt Reference Proteomes, we decided to study the bacteria with the highest number of FA-like proteins per proteome size and investigate their pathogenicity, listed in Table 1. Several of them are known pathogens, like the food-born pathogen *Listeria monocytogenes* [23]. Other strains are not yet well characterized, but are known to be able to cause infections, for example *Sneathia amnii*, which can cause urethritis and can affect pregnancy [24]. Whereas *Veillonella* species belong to the normal flora in the gastrointestinal tract, mouth and vaginal tract and are rarely related to infections [25]. We could also detect FA-like proteins for *Staphylococcus epidermidis* or *Bifidobacterium breve*, which are known to function beneficially [26, 27]. Thus presence of FA-like proteins in a proteome might indicate possible pathogenicity, but it does not solely determine it.

**Table 1:** Bacterial strains with the highest fraction of FA-like proteins per proteome size.

| Organism | Proteome ID (UniProt) | Proteome size | FA-like proteins | Fraction | Pathogenicity |
| --- | --- | --- | --- | --- | --- |
| *Streptobacillus moniliformis* (strain ATCC 14647 / DSM 12112 / NCTC 10651 / 9901) | UP000002072 | 1431 | 21 | 0.0147 | Yes |
| *Histophilus somni* (strain 2336) | UP000008543 | 1971 | 12 | 0.0061 | Yes |
| *Sneathia amnii* | UP000033103 | 1182 | 6 | 0.0051 | Yes |
| *Listeria monocytogenes* serovar 1/2a (strain ATCC BAA-679 / EGD-e) | UP000000817 | 2844 | 14 | 0.0049 | Yes |
| *Myroides indicus* | UP000295215 | 2689 | 13 | 0.0048 | Opport. |
| *Basilea psittacipulmonis* DSM 24701 | UP000028945 | 1452 | 7 | 0.0048 | Unknown |
| *Veillonella criceti* | UP000255367 | 1926 | 9 | 0.0047 | Unknown |
| *Taylorella asinigenitalis* (strain MCE3) | UP000009284 | 1523 | 7 | 0.0046 | Unknown |
| *Haemophilus parasuis* serovar 5 (strain SH0165) | UP000006743 | 2002 | 9 | 0.0045 | Yes |
| *Streptococcus sanguinis* (strain SK36) | UP000002148 | 2269 | 10 | 0.0044 | Opport. |
| *Mycoplasma* sp. CAG:877 | UP000018280 | 1407 | 6 | 0.0043 | Unknown |
| *Luteimonas* sp. H23 | UP000308707 | 3308 | 14 | 0.0042 | Unknown |
| *Actinobacillus seminis* | UP000215738 | 2028 | 8 | 0.0039 | Yes |
| *Weissella ceti* | UP000029079 | 1338 | 5 | 0.0037 | Yes |
| *Clostridium* sp. CAG:354 | UP000018313 | 1707 | 6 | 0.0035 | Unknown |
| *Pseudoscardovia suis* | UP000216454 | 1732 | 6 | 0.0035 | Unknown |
| *Myroides guanonis* | UP000243887 | 2674 | 9 | 0.0034 | Unknown |
| *Megasphaera* sp. UPII 135-E | UP000004137 | 1512 | 5 | 0.0033 | Unknown |
| Firmicutes bacterium CAG:321 | UP000017979 | 1220 | 4 | 0.0033 | Unknown |
| *Veillonella* sp. oral taxon 780 str. F0422 | UP000010295 | 1585 | 5 | 0.0032 | Unknown |

Unknown: no study investigating pathogenicity found for this organism or specific species, or species not given.
Opport.: opportunistic pathogen.

## Discussion

In this paper we aimed to comprehensively identify FA-like proteins and show their characteristics with respect to their domain composition, domain architecture and taxonomic distribution. Our approach based on using Pfam Hidden Markov Models (HMMs) of the known adhesive and stalk domains has resulted in the discovery of many novel FA-like proteins from a wide range of bacterial genera. These proteins include new putative binding proteins in well studied pathogens like *Listeria monocytogenes*, in poorly characterised pathogens, like *Sneathia amnii*, but also in commensal bacteria, like *Bifidobacterium breve*, indicating that fibrillar adhesins can be involved in pathogenesis, but also in commensal interactions.

Our results show the various different adhesive and stalk domain combinations that exist in nature and also how frequently they are found. The information can facilitate the identification of new fibrillar adhesins. For instance, when analysing a new FA-like protein, a potential annotated adhesive domain can infer the existence of a specific stalk domain and vice versa. Domain grammar approaches have shown that a knowledge of the domain grammar of proteins can enhance sequence similarity searching [28, 29]. For a representative analysis of the phylogenetic distribution of FA-like proteins we decided to use the UniProt Reference Proteomes sequences only. When using the whole UniProtKB database, more FA-like proteins and more adhesive and stalk domain combinations can be found (supplementary figure S4). However, using the whole UniProtKB would lead to a lot of redundancy that would cause significant biases in our observations. The observation that several adhesive domains are found in combination with different stalk domains raises the question, whether adhesive domains can function with any arbitrary stalk. We suggest that this would be an interesting question for experimental studies to address.

One of the more striking results with respect to domain grammar was the strength of the presence of adhesive domains to be found at the N-terminus of Firmicutes FA-like proteins. This preference is largely due to the presence of a C-terminal sortase motif in these proteins. However, even when we investigate Firmicutes proteins without a sortase anchor we still see this N-terminal preference for adhesive domains. This seems likely due to our weak ability to detect short anchoring motifs. In nearly half of the detected FA-like proteins no known anchor was found, suggesting novel anchor domains or motifs may yet exist to be discovered.

The investigation of the taxonomic distribution of adhesive and stalk domains show a wide range of different domains in Gram positive bacteria. One reason for this variety might be that the majority of fibrillar adhesins already studied belong to Firmicutes genera, as for example *S. aureus* [30], and subsequently a large number of adhesive domains on our list were first discovered in Firmicutes. Most of the FA-like proteins in the Gram negative Proteobacteria are composed of YadA_head and YadA_stalk domains. The lack of information about fibrillar adhesins in Gram negative bacteria is one possible reason for the low diversity of FA-like protein hits in Proteobacteria. Another possibility is that Gram negative bacteria might predominantly use pili for the adhesion to host cells and therefore possess less fibrillar adhesins. The taxonomic investigation furthermore demonstrates that there are domains which are found to be common between widely diverged phyla such as Firmicutes and Actinobacteria. But it also shows that there are domains which are only found in a specific phylum such as Intimin_C. In the case of Intimin_C we know from its structure that it is distantly related to other C-type lectin domains. In these cases of phylogenetically restricted domains it will be interesting to understand to what extent they have evolved novel binding specificities.

In our results we describe that the adhesive domain SSURE is not found in combination with other known stalk domains, and that it is found on average in 3 copies per protein. We additionally know that SSURE is tandemly repeated. This leads to the hypothesis that some adhesive domains might be able to function as the stalk itself. Several adhesive domains are indeed found repeated. A reason for repeating adhesive domains could be to increase the avidity of binding to the host cells. Thus duplications of these domains may lead to both extending the stalk length and increasing binding efficiency. In particular, when the host ligand is a collagen, the adhesive domain might bind to different positions simultaneously on the host ligand. The adhesive domain Collagen_bind binds collagen with the very specific collagen hug mechanism which requires the supra-domain of Big_8 domain N-terminal to the Collagen_bind domain. Therefore, it would be interesting to find out why Collagen_bind occurs in repeats and whether all repeats contribute to the adhesion process.

The greatest limitation of our FA-like protein discovery process is that it requires a protein to contain a combination of known stalk and adhesive domains. There are undoubtedly numerous fibrillar adhesive proteins missed by our approach. There are several reasons for these missed proteins. Firstly, there may be FA-like proteins that are missing domain annotations for either the stalk or adhesive domain or both. This could be because there is no domain family represented within Pfam currently. A second possibility is that the current Pfam model misses true examples of the stalk or adhesive domain. And thirdly, simply that there are adhesive and stalk domains we did not yet identify. An estimation of the completeness of the stalk domain list for known adhesive domains is given in supplementary table S4. Nevertheless, the information about fibrillar adhesins gained in this study will be important for further improvements in future identification of these proteins.

## Conclusions

This study presents to our knowledge the most comprehensive characterisation of fibrillar adhesins to date. We have used a protein domain based screen to identify over 3,000 fibrillar adhesin-like proteins and characterised their domain content, architecture and taxonomic distribution. The results underline the complexity of fibrillar adhesins with highly variable domain organisations as well as with various different domain combinations found in nature. The detected FA-like proteins are widespread across the bacterial taxonomic tree in pathogenic as well as in non-pathogenic bacteria. The set of FA-like proteins and associated stalk and adhesive domains identified in this study set a foundation stone to discover further fibrillar adhesins.

## Methods

### Collection of Stalk, Adhesive and Anchor domains

We searched in the literature for experimentally confirmed bacterial adhesins, for example by a keyword search for ‘adhesins’ in PubMed. We analysed the domain architecture of the relevant proteins and identified the adhesive domains by searching for the binding regions. We also identified numerous adhesive domains through investigation of structures in the Protein Data Bank. In each case we identified the relevant entries in the Pfam database and recorded the relevant identifiers.

Stalk domains were detected by searching for tandem repeat domains in the Pfam database, whereby repeats are counted as tandem repeats when the sequence separation between two domains from the same Pfam family is shorter than 35 residues. The further putative stalk domain families were checked manually in Pfam to ensure their description, species distribution and domain architecture matched the stalk domain definition. Domain families found only in eukaryotes, and non-surface and enzymatic proteins, were excluded.

Tandem sequence repeats were identified in proteins without known Pfam annotations with the T-REKS tool using 70% sequence identity and length between 50-200 residues as parameters [31]. Six new Pfam families were built (Pfam: PF19403 - PF19408) by first clustering sequence repeats with BLAST and then iterating alignments of the largest clusters [32].

### Identification of Fibrillar Adhesin-like proteins

In this study all bacterial proteins from the UniProt Reference Proteomes (release 2020_03) were used [8]. The 7,581 bacterial reference proteomes encompass 28,086,379 proteins. We searched with the identified adhesive and stalk Pfam domain HMMs against the bacterial protein sequences using the HMMER tool (version 3.1b2) with the gathering (GA) threshold option [33]. Overlapping annotations were excluded based on the hmmsearch annotation score. We identified proteins with at least one known adhesive domain (supplementary list S2) and one known stalk domain (supplementary list S1) and constructed a fasta sequence database for the detected proteins. The matches from this query were considered as FA-like proteins and were used for the majority of analyses. For comparison, FA-like proteins were also searched for in the whole UniProtKB using the same approach.

### Characterization of Fibrillar Adhesin-like proteins

The number of stalk domains per protein in combination with one adhesive domain were counted, regardless of whether the stalk domains belonged to the same Pfam family or not. The frequency of adhesive and stalk domains as well as the combinations of adhesive and stalk domains and different stalk domains found on the same FA-like proteins were counted, whereby each domain was only counted once even if multiple domains of the same family were annotated on the protein.

For the generation of the density plots representing the domain positions within the proteins, the protein length and the envelope domain location information were used from the HMMER search output file. The fractional start and end positions for the domains were calculated. Furthermore, sortase motifs were identified by regular expression in the C-terminal 50 amino acids of the FA-like protein sequences. For the analysis of the domain architecture between the different phyla, the taxonomic phyla were mapped to the UniProt protein identifier by using Retrieve/ID mapping from the UniProt website (https://www.uniprot.org/uploadlists/).

### Identification of cell anchoring mechanisms of FA-like proteins

To find out how FA-like proteins are attached to the bacterial cell, HMMs of known anchor domains were downloaded from Pfam. We used the following models: Gram_pos_anchor (Pfam: PF00746), SLH (Pfam: PF00395), CHU_C (Pfam: PF13585), Por_Secre_Tail (Pfam: PF18962), LysM (Pfam: PF01476), Choline_bind_1 (Pfam: PF01473), Choline_bind_2 (Pfam: PF19085), Choline_bind_3 (Pfam: PF19127), YadA_anchor (Pfam: PF03895) IAT_beta (Pfam: PF11924), GW (Pfam: PF13457), OMP_B-brl (Pfam: PF13505), PG_binding_1 (Pfam: PF01471), PG_binding_2 (Pfam: PF08823) and PG_binding_3 (Pfam: PF09374) [34]. By HMMER (version 3.2.1) hmmsearch using each profile’s Gathering (GA) threshold the FA-like protein sequences were searched for the above listed anchor domains [33]. To improve the search sensitivity for Gram positive anchor motifs, we carried out additional regular expression searches for the canonical sequence motifs LPxTA and LPxTG as well as for the alternative anchor motifs NPxTG, LAxTG, NPQTN and LPxGA. To be considered a match we also required these regular expression matches to occur within the C-terminal 50 amino acids of the sequence. If these sortase motif protein hits were missed by the hmmsearch for the Gram_pos_anchor, the identifiers were added and are displayed together as sortase motifs in the bar plots. It was shown that RTX adhesins can be inserted into a beta barrel of an outer membrane pore with its N-terminus [15]. Shuaiqi Guo *et al*. (2017) aligned the N-terminus of ten RTX adhesins of different Gram negative bacteria and showed that an alpha-helical element (‘xxIQxAIAA’), followed by a short coil motif (‘GxDPT’) and by the LapG protease cleavage motif (‘uAAG’, where the u is a threonine or proline residue) are well aligned in these proteins [15]. By using their ten representative RTX adhesin sequences the alignment was reproduced and an HMM was built, which yielded 37 hits in all 3388 FA-like protein sequences. But even in known RTX adhesins, like MhLap of *Marinobacter hydrocarbonoclasticus* (UniProt: H8W6K8), the described secondary structure motifs were not found so we only searched for the protease cleavage motif ‘AAG’ within the first 150 amino acids in the FA-like protein sequences. To increase the probability that the found hits are indeed RTX adhesins, we additionally searched for the HemolysinCabind domain, which is characteristic for the RTX adhesins. In total 62 LapG cleavage motifs and HemolysinCabind domains could be found together in a single protein. In 31 of these 62 FA-like protein hits the short coil motif ‘GxDPT’ could be found, whereas the alpha-helical element motif ‘xxIQxAIAA’ was only found in two of these proteins. For comparison, the PSORTb webserver (version 3.0.2) was used to predict the localization of the 3,388 FA-like proteins [18].

### Taxonomic analysis

For taxonomic analysis, additional to the phyla, the genera were also mapped by using Retrieve/ID mapping from the UniProt website. The 16s rRNA sequences of representative bacterial genera and of genera with a minimum of 5 detected FA-like proteins were aligned in JalView using Muscle with the default settings [35, 36]. Afterwards a Neighbour Joining based tree was calculated with JalView by using the BLOSUM62 matrix for measuring the distance of the alignments [35]. For the visualisation of the tree iTOL was used [37]. For each genera the adhesive and stalk domains of FA-like proteins were counted and mapped to the tree. Again, stalk domain repeats on the same FA-like protein were counted as one occurrence.

### Searching for adhesive and stalk repeats

Protein sequences containing known adhesive domains were downloaded from Pfam version [34]. Proteins were further filtered to be from bacterial species and a minimum of 500 amino acids long, and searched against Pfam using the HMMER tool (version 3.2.1) with the gathering (GA) threshold option [33]. Overlapping annotations were again excluded based on the hmmsearch annotation score. The number of stalk and adhesive domain annotations per protein were counted and the average was calculated per adhesive domain family.

### Investigation of completeness of stalk domain list

The HMMER search results of the long bacterial proteins with known adhesive domains against the Pfam database from the section ‘Searching for adhesive and stalk repeats’ were used again. Proteins with known stalk domains were counted as well as proteins without any Pfam domain annotation, except for the anchor domains listed in supplementary table S3, the signal domains YSIRK_signal (Pfam: PF04650), ESPR (Pfam: PF13018) and the TAT_signal (Pfam: PF10518) and the domains in known supra domains.

## Supplementary

**Table S1:** Stalk domains identified and used in this study

| Name | Pfam ID | Pfam Clan | Clan ID |
| --- | --- | --- | --- |
| Rib | PF08428 | E-set | CL0159 |
| Big_1 | PF02369 | E-set | CL0159 |
| Big_2 | PF02368 | E-set | CL0159 |
| Big_3 | PF07523 | E-set | CL0159 |
| Big_3_5 | PF16640 | E-set | CL0159 |
| Big_6 | PF17936 | E-set | CL0159 |
| Big_9 | PF17963 | E-set | CL0159 |
| DUF5011 | PF16403 | E-set | CL0159 |
| DUF11 | PF01345 | E-set | CL0159 |
| PKD | PF00801 | E-set | CL0159 |
| He_PIG | PF05345 | E-set | CL0159 |
| Cadherin_4 | PF17803 | E-set | CL0159 |
| Cadherin_5 | PF17892 | E-set | CL0159 |
| CARDB | PF07705 | E-set | CL0159 |
| Calx-beta | PF03160 | E-set | CL0159 |
| TIG | PF01833 | E-set | CL0159 |
| Cadherin | PF00028 | E-set | CL0159 |
| fn3 | PF00041 | E-set | CL0159 |
| Big_13 | PF19077 | E-set | CL0159 |
| Big_12 | PF19078 | E-set | CL0159 |
| Big_11 | PF18200 | E-set | CL0159 |
| Big_5 | PF13205 | E-set | CL0159 |
| Big_3_2 | PF12245 | E-set | CL0159 |
| HYR | PF02494 | E-set | CL0159 |
| InIK_D3 | PF18981 | E-set | CL0159 |
| PKD_4 | PF18911 | E-set | CL0159 |
| Cadherin_3 | PF16184 | E-set | CL0159 |
| PKD_5 | PF19406 | E-set | CL0159 |
| PKD_6 | PF19408 | E-set | CL0159 |
| SHIRT | PF18655 | Ubiquitin | CL0072 |
| Flg_new | PF09479 | Ubiquitin | CL0072 |
| MucBP | PF06458 | Ubiquitin | CL0072 |
| SSSPR-51 | PF18877 | Ubiquitin | CL0072 |
| MucBP_2 | PF17965 | Ubiquitin | CL0072 |
| Flg_new_2 | PF18998 | Ubiquitin | CL0072 |
| DUF1542 | PF07564 | B_GA | CL0598 |
| GA | PF01468 | B_GA | CL0598 |
| FIVAR | PF07554 | B_GA | CL0598 |
| MBG | PF17883 | MBG | CL0682 |
| MBG_2 | PF18676 | MBG | CL0682 |
| MBG_3 | PF18887 | MBG | CL0682 |
| FctA | PF12892 | Transthyretin | CL0287 |
| SpaA | PF17802 | Transthyretin | CL0287 |
| Cna_B | PF05738 | Transthyretin | CL0287 |
| SdrD_B | PF17210 | Transthyretin | CL0287 |
| CarboxypepD_reg | PF13620 | Transthyretin | CL0287 |
| SpaA_2 | PF19403 | Transthyretin | CL0287 |
| DUF5979 | PF19407 | Transthyretin | CL0287 |
| G5 | PF07501 | G5 | CL0593 |
| LVIVD | PF08309 | Beta_propeller | CL0186 |
| DUF5122 | PF17164 | Beta_propeller | CL0186 |
| SBBP | PF06739 | Beta_propeller | CL0186 |
| Ig_7 | PF19081 | Ig | CL0011 |
| I-set | PF07679 | Ig | CL0011 |
| TSP3_bac | PF18884 | TSP3 | CL0689 |
| DUF285 | PF03382 | LRR | CL0022 |
| SlpA | PF03217 | No clan | No clan |
| Agl_II_C2 | PF17998 | No clan | No clan |
| Collagen | PF01391 | No clan | No clan |
| Fn_bind | PF02986 | No clan | No clan |
| CshA_repeat | PF19076 | No clan | No clan |
| Antigen_C | PF16364 | No clan | No clan |
| TQ | PF18202 | No clan | No clan |
| YadA_stalk | PF05662 | No clan | No clan |
| Endotoxin_C2 | PF18449 | No clan | No clan |
| Trp_ring | PF18669 | No clan | No clan |
| Strep_SA_rep | PF06696 | No clan | No clan |
| DUF5801 | PF19116 | No clan | No clan |
| DUF1079 | PF06435 | No clan | No clan |
| Octapeptide | PF03373 | No clan | No clan |
| DUF5776 | PF19087 | No clan | No clan |
| QPE | PF18874 | No clan | No clan |
| CFSR | PF19079 | No clan | No clan |
| DUF5977 | PF19404 | No clan | No clan |
| DUF5978 | PF19405 | No clan | No clan |

**Table S2:**
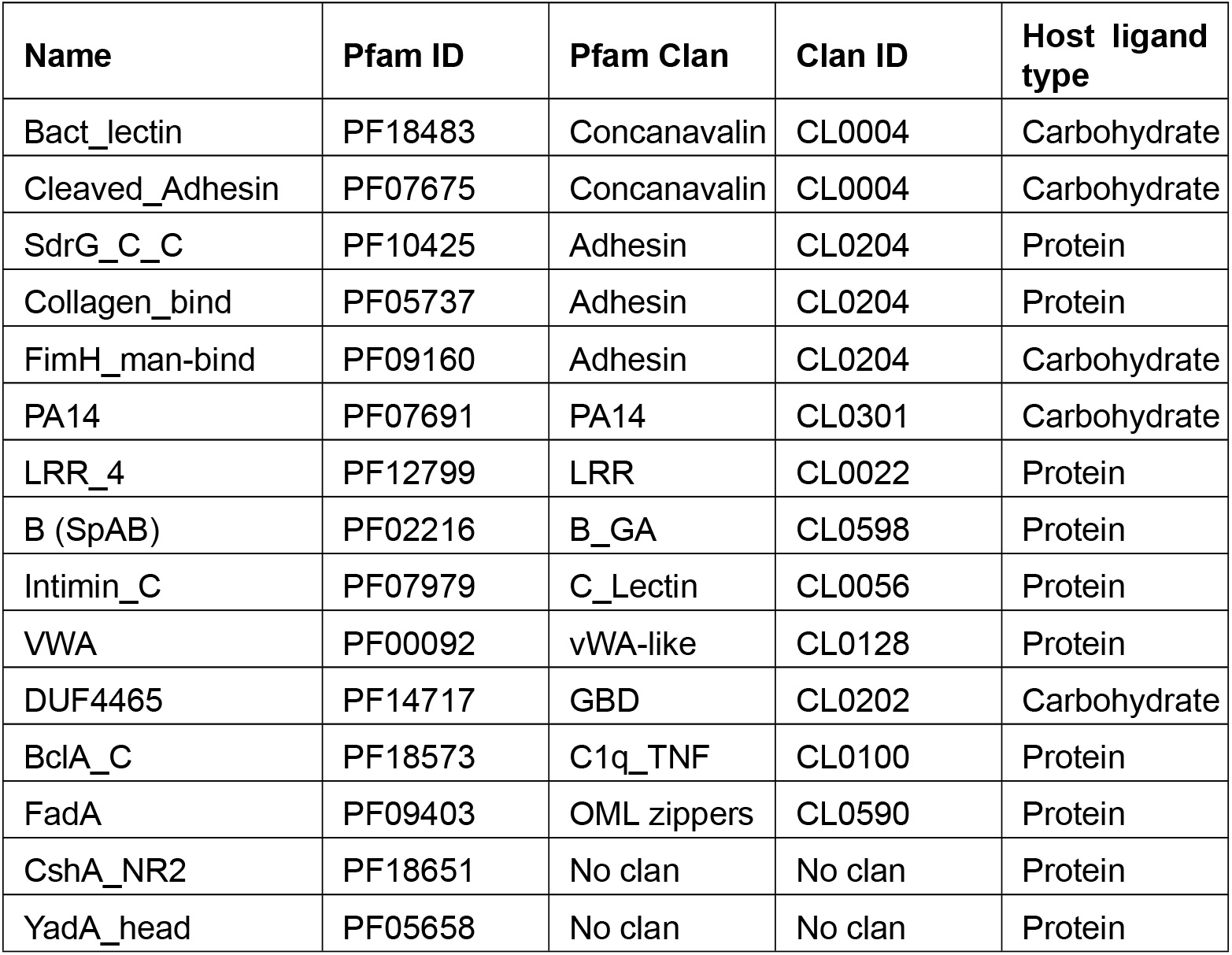
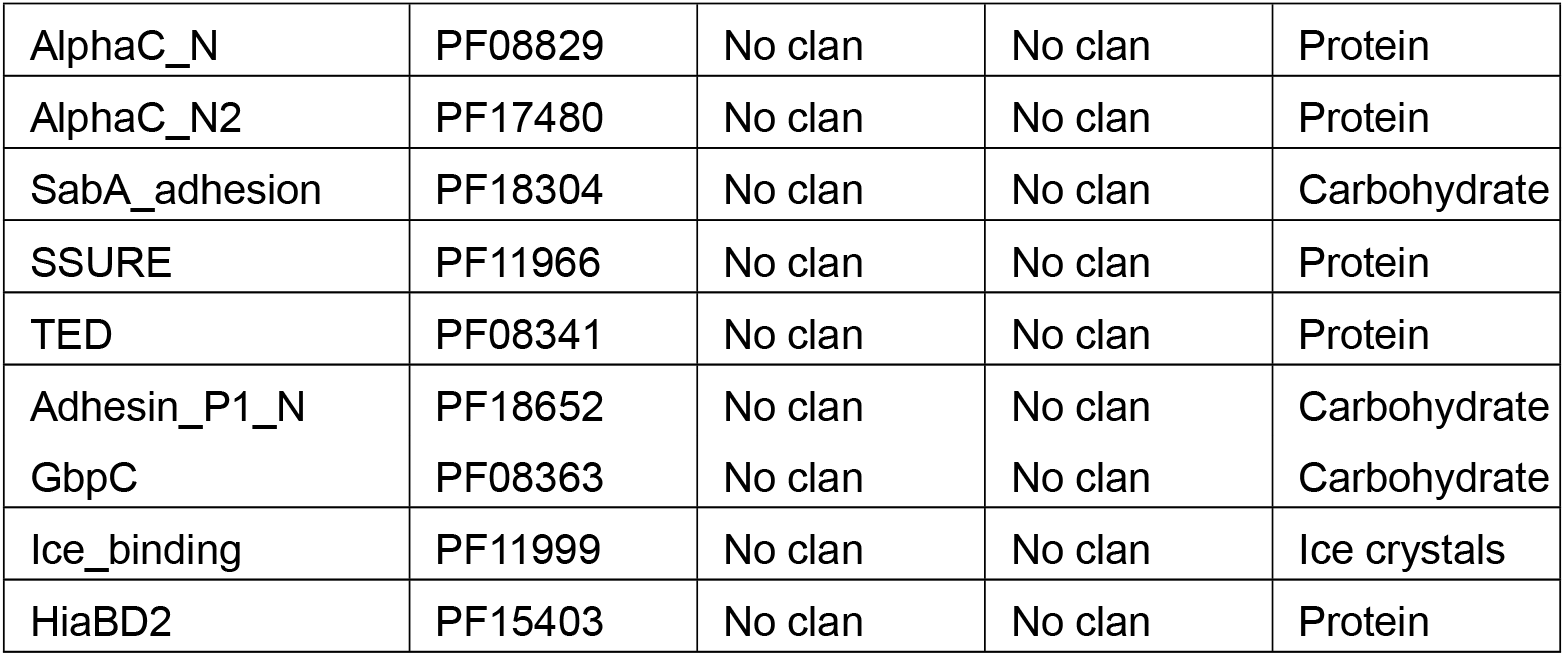
Adhesive domains used in this study

| Name | Pfam ID | Pfam Clan | Clan ID | Host ligand type |
| --- | --- | --- | --- | --- |
| Bact_lectin | PF18483 | Concanavalin | CL0004 | Carbohydrate |
| Cleaved_Adhesin | PF07675 | Concanavalin | CL0004 | Carbohydrate |
| SdrG_C_C | PF10425 | Adhesin | CL0204 | Protein |
| Collagen_bind | PF05737 | Adhesin | CL0204 | Protein |
| FimH_man-bind | PF09160 | Adhesin | CL0204 | Carbohydrate |
| PA14 | PF07691 | PA14 | CL0301 | Carbohydrate |
| LRR_4 | PF12799 | LRR | CL0022 | Protein |
| B (SpAB) | PF02216 | B_GA | CL0598 | Protein |
| Intimin_C | PF07979 | C_Lectin | CL0056 | Protein |
| VWA | PF00092 | vWA-like | CL0128 | Protein |
| DUF4465 | PF14717 | GBD | CL0202 | Carbohydrate |
| BclA_C | PF18573 | C1q_TNF | CL0100 | Protein |
| FadA | PF09403 | OML zippers | CL0590 | Protein |
| CshA_NR2 | PF18651 | No clan | No clan | Protein |
| YadA_head | PF05658 | No clan | No clan | Protein |
| AlphaC_N | PF08829 | No clan | No clan | Protein |
| AlphaC_N2 | PF17480 | No clan | No clan | Protein |
| SabA_adhesion | PF18304 | No clan | No clan | Carbohydrate |
| SSURE | PF11966 | No clan | No clan | Protein |
| TED | PF08341 | No clan | No clan | Protein |
| Adhesin_P1_N | PF18652 | No clan | No clan | Carbohydrate |
| GbpC | PF08363 | No clan | No clan | Carbohydrate |
| Ice_binding | PF11999 | No clan | No clan | Ice crystals |
| HiaBD2 | PF15403 | No clan | No clan | Protein |

**Table S3:**
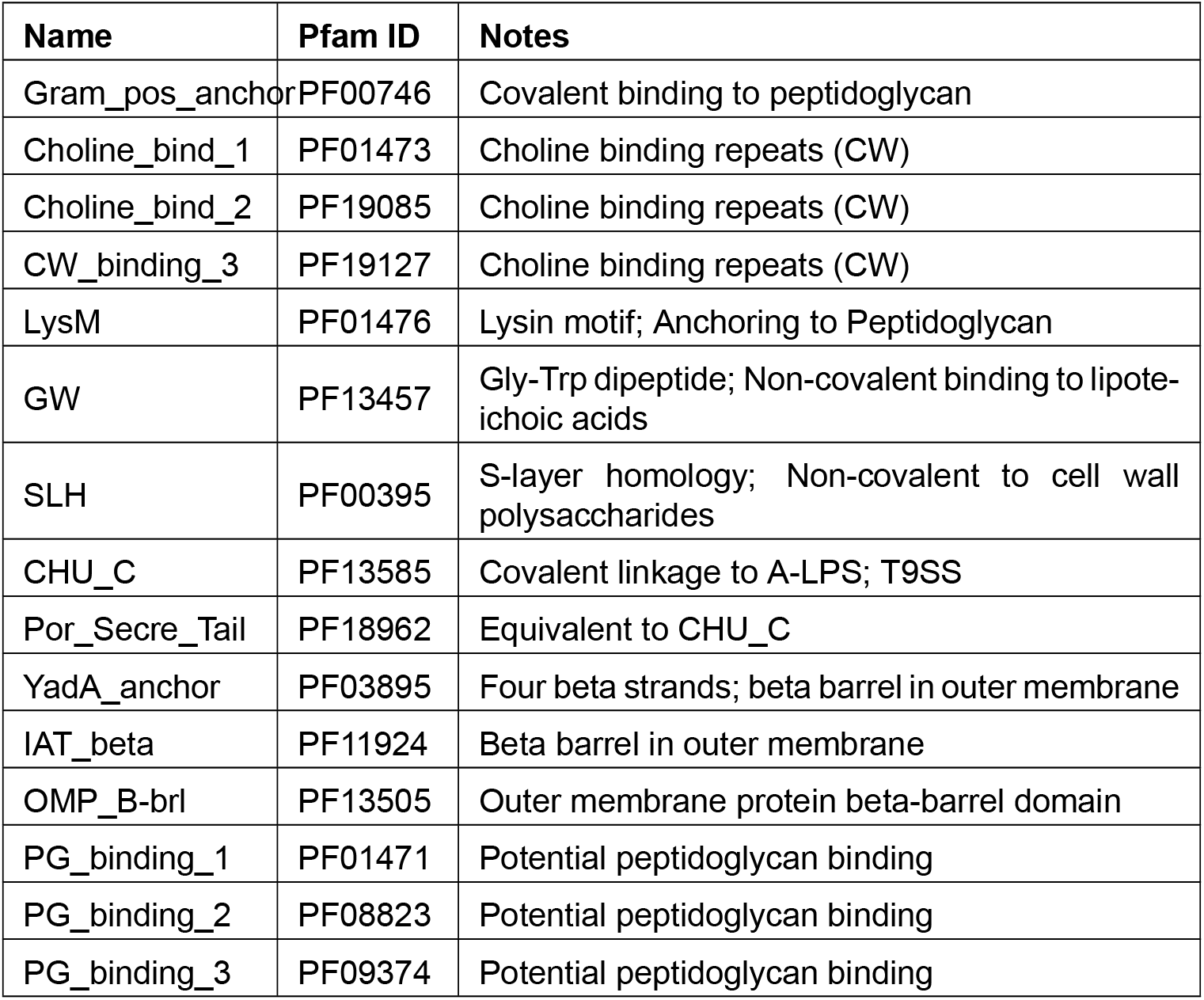
Anchor domains used in this study

**Table S4:** Investigation of completeness of the stalk domain list. Bacterial proteins in Pfam with a length of minimum 500 residues were counted per adhesive domain and listed in the second column ‘Long proteins’. The coverage of long proteins with known stalk domains are counted in the third column ‘With known stalk’. The fourth column represents the number of long proteins, without any Pfam annotation and therefore proteins with potential uncharacterized stalk domains.

| **Name** | **Long proteins** | **With known stalk** | **W/o annotations** |
| --- | --- | --- | --- |
| Bact_lectin | 396 | 165 | 129 |
| Cleaved_Adhesin | 206 | 103 | 75 |
| SdrG_C_C | 51 | 38 | 13 |
| Collagen_bind | 399 | 365 | 33 |
| PA14 | 2915 | 303 | 318 |
| LRR_4 | 527 | 88 | 133 |
| B (SpAB) | 1 | 1 | 0 |
| Intimin_C | 2 | 2 | 0 |
| VWA | 4563 | 490 | 775 |
| DUF4465 | 39 | 3 | 15 |
| BclA_C | 28 | 25 | 3 |
| CshA_NR2 | 28 | 25 | 3 |
| YadA_head | 1074 | 877 | 150 |
| AlphaC_N | 1 | 1 | 0 |
| AlphaC_N2 | 1 | 1 | 0 |
| SabA_adhesion | 6 | 0 | 0 |
| SSURE | 11 | 0 | 11 |
| TED | 266 | 243 | 19 |
| Adhesin_P1_N | 36 | 26 | 10 |
| GbpC | 65 | 31 | 33 |
| Ice_binding | 198 | 133 | 41 |

**Figure S1:**
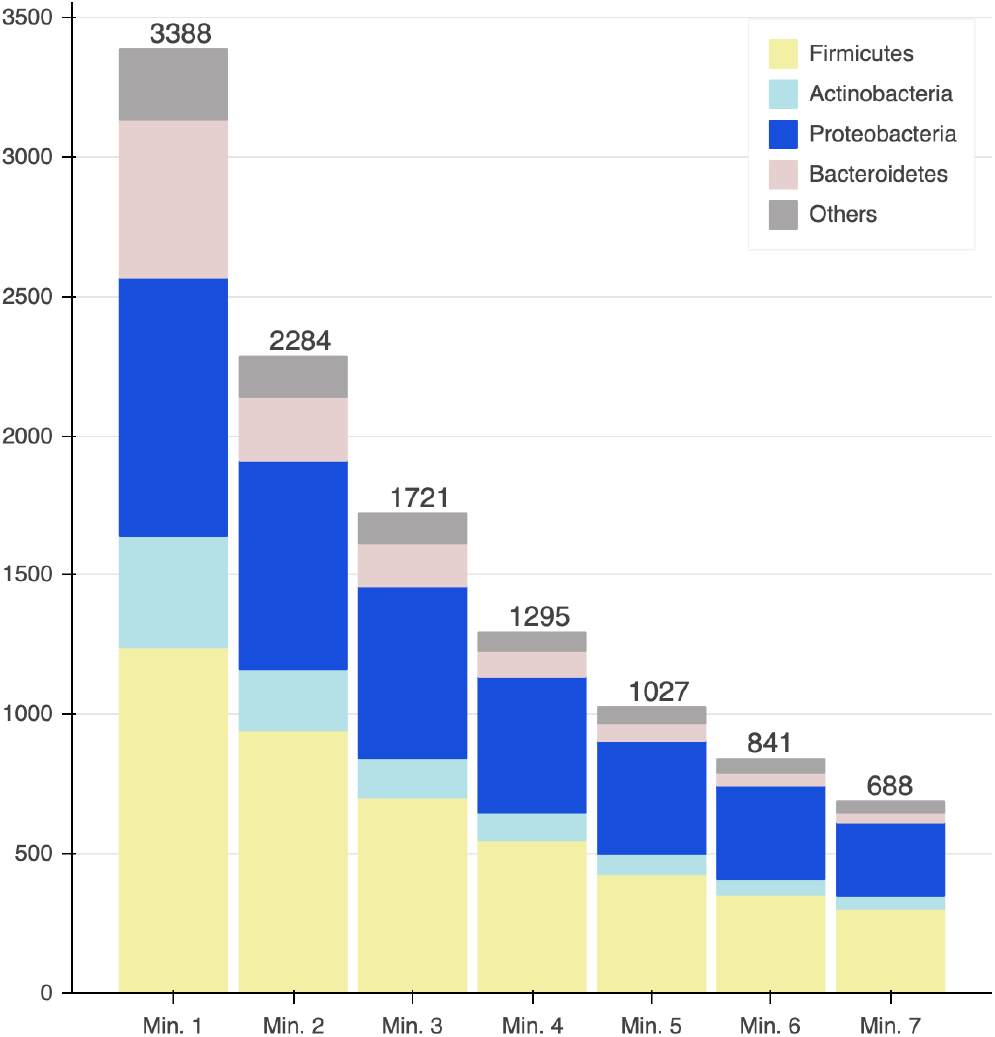
Number of stalk domains per FA-like protein per phyla: Graph showing the number of FA-like proteins identified in our screen depending on the number of stalk domains required. It is split between the most prevalent phyla in the dataset.

**Figure S2:**
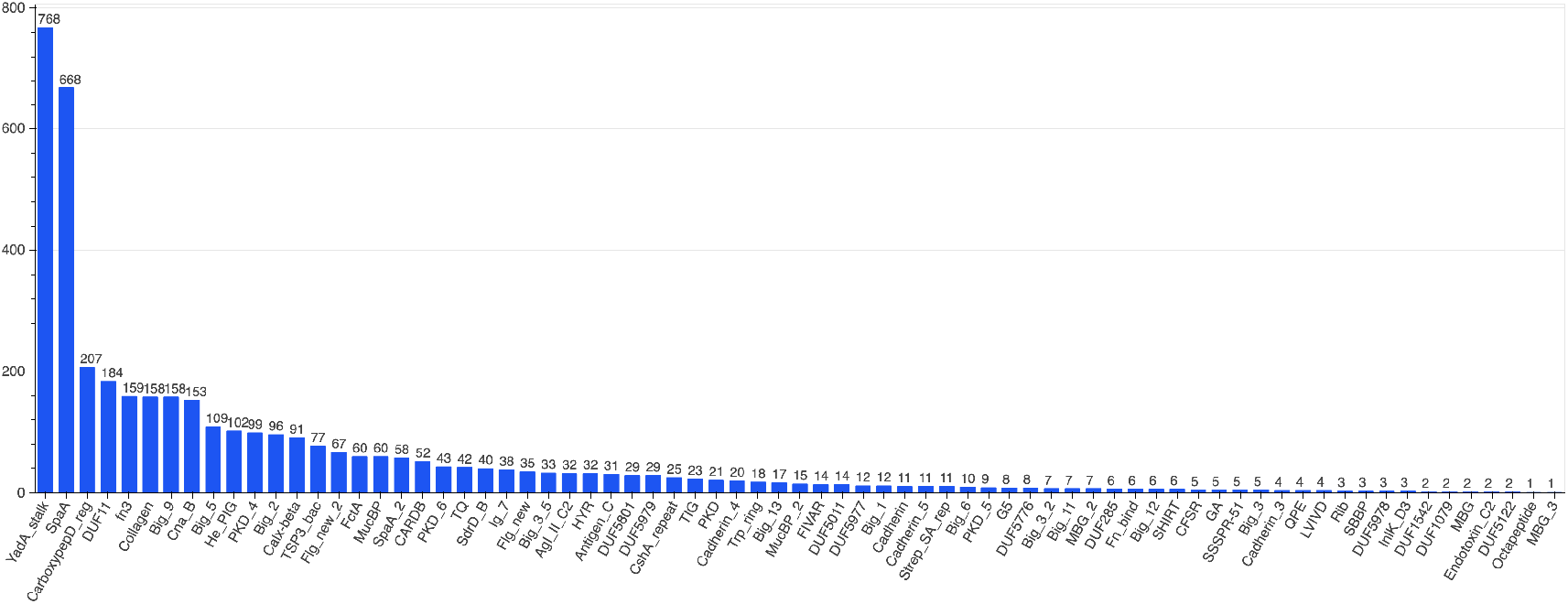
Frequency of stalk domains in combination with adhesive domains: Number of FA-like proteins in which the stalk domains were detected.

**Figure S3:**
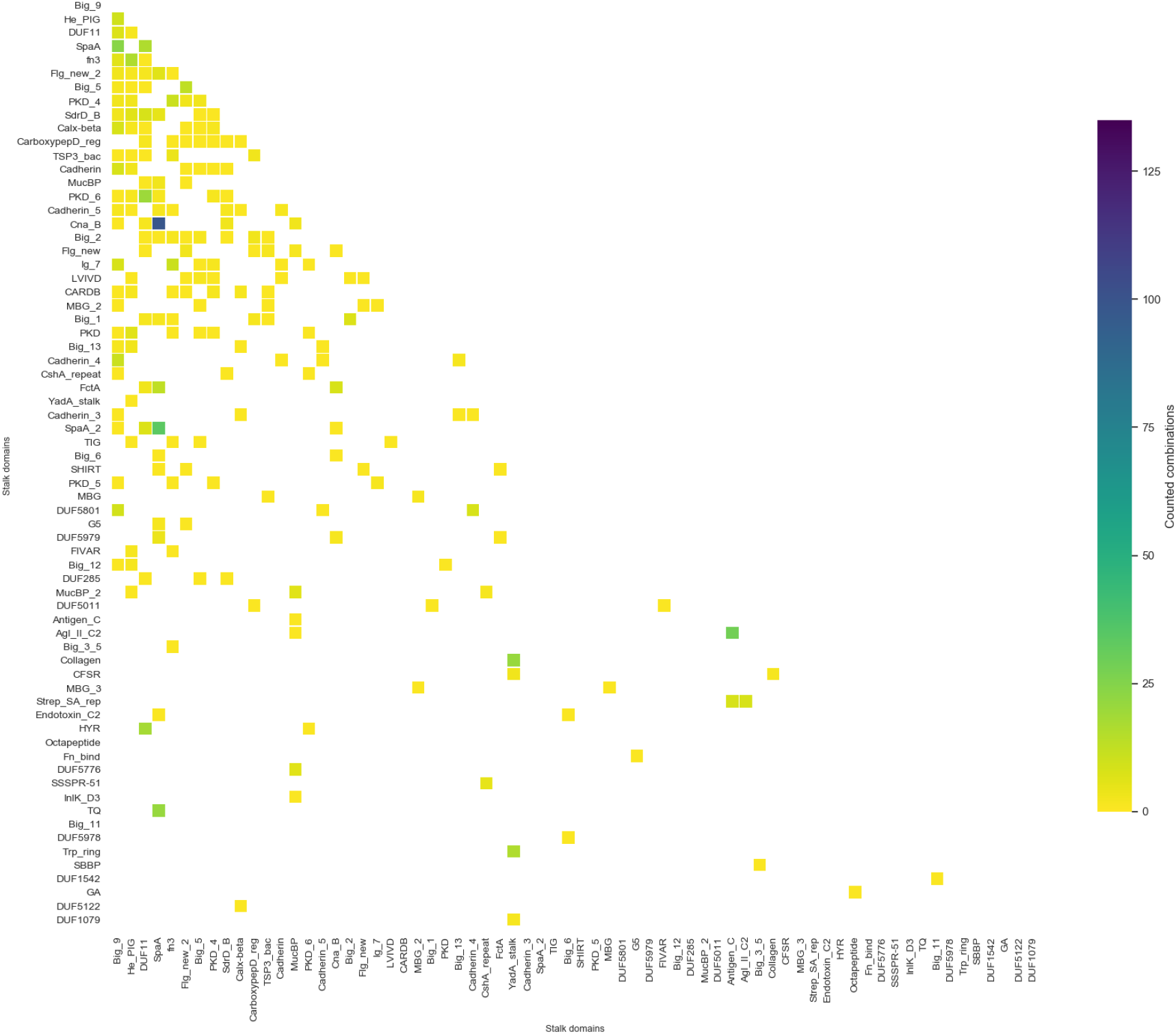
Domain combinations of stalk domains: Heatmap showing the co-occurrence of stalk domains on the same FA-like protein.

**Figure S4:**
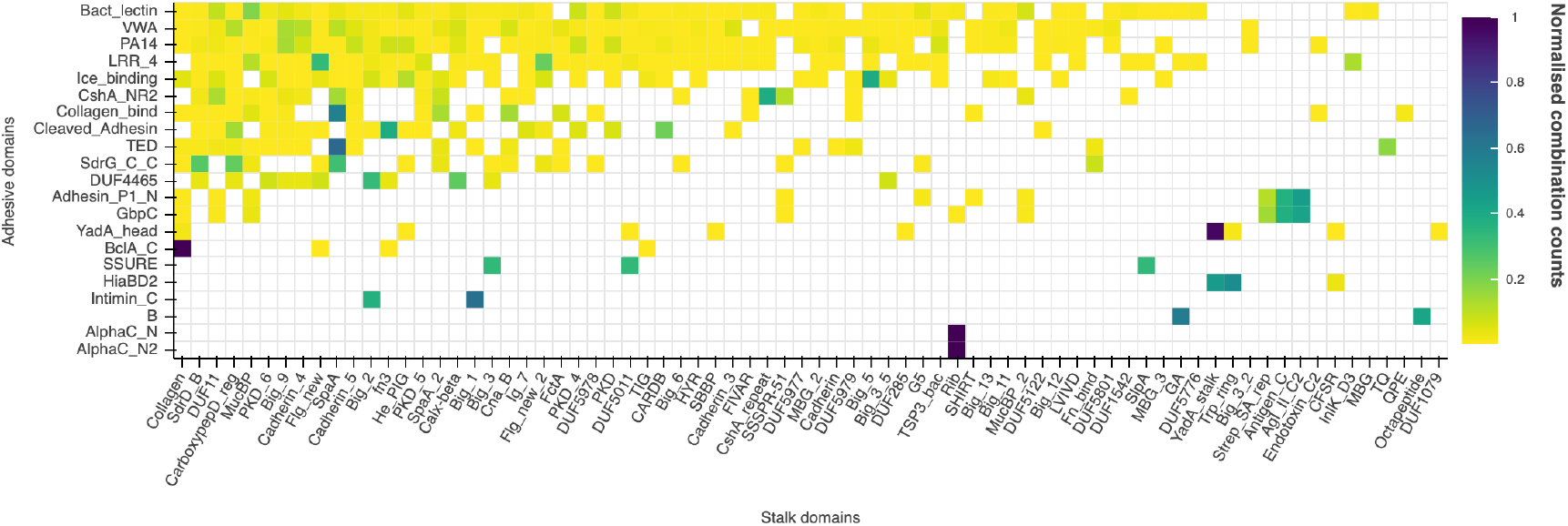
Domain combinations of adhesive and stalk domains found in UniProtKB: Compared to figure 3, this heatmap is based on detected FA-like proteins not only found in the reference proteomes, but the whole UniProtKB.

**Figure S5:**
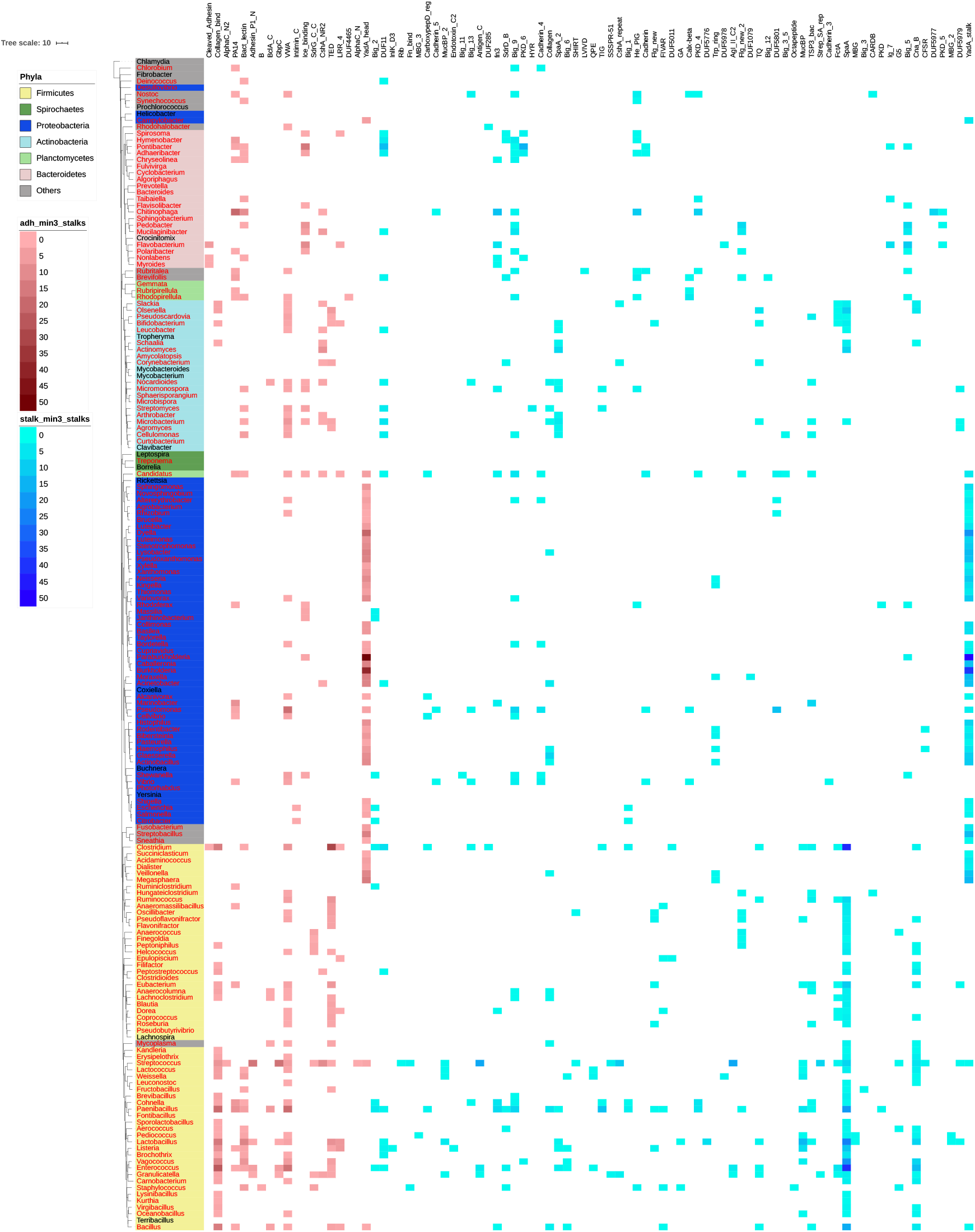
Overview of taxonomic hits for proteins with at least one adhesive and at least three known stalk domains: This taxonomic tree shows only these FA-like proteins were additional to the adhesive domain minimal 3 known stalk domains could be annotated.

## Declarations

### Ethics approval and consent to participate

Not applicable.

### Consent for publication

Not applicable.

### Availability of data and materials

The FA-like protein dataset generated during the current study is available in the GitHub repository https://github.com/VivianMonzon/FA-like_proteins.

### Authors’ contributions

AB conceived and supervised the study and was a major contributor in writing the manuscript. VM conducted the identification and characterization of FA-like proteins and wrote the manuscript. AL conducted the screen for stalk domains and was a major contributor in writing the manuscript. All authors read, revised and approved the final manuscript.

## Acknowledgement

Not applicable.

## Authors’ information (optional)

